# Systematic identification of yeast mutants with increased rates of cell death reveals rapid stochastic necrosis associated with cell division

**DOI:** 10.1101/2021.10.20.465133

**Authors:** Alexander I. Alexandrov, Erika V. Grosfeld, Olga V. Mitkevich, Victoria A. Bidyuk, Arina V. Nostaeva, Igor V. Kukhtevich, Robert Schneider, Evgeniy S. Shilov, Vitaliy V. Kushnirov, Sergey E. Dmitriev, Vadim N. Gladyshev

## Abstract

Cell death plays a major role in development, pathology and aging and can be triggered by various types of acute stimuli which arrest cell growth. However, little is known about chronic cell death in the context of continuing cell division. Here, we performed a genome-wide search for mutants with this type of death in dividing baker’s yeast by assaying staining with phloxine B, which accumulates in dead cells. This screen yielded 83 essential and 43 non-essential gene mutants. Three contrasting types of spatial distribution of dead cells in colonies were observed, which corresponded to gene ontology enrichment for (i) DNA replication and repair, RNA processing, chromatin organization, and nuclear transport; (ii) mitosis and cytokinesis; and (iii) vesicular transport and glycosylation/cell wall homeostasis. To study dynamics of cell death in these mutants, we developed methods for analyzing the death of newborn cells (DON) and cell death in real time using microfluidics-based microscopy. These revealed rapid stochastic necrosis during bud generation or cytokinesis without prior division arrest. Increased death during division was associated with common sensitivity to plasma membrane and cell-wall perturbing agents, and could be mitigated by neutral pH stabilization of the medium. This suggests a common downstream type of cell death caused by a wide range of genetic perturbations.

## Introduction

The cell is a complex agglomerate of components and systems that act in concert to achieve sustainable replication of either itself or, for a multicellular organism, of the organism’s germ line. Cellular machinery is largely made up of proteins, some of which are more dispensable than others (1). The absence of indispensable (essential) proteins makes cells inviable, precludes them from replicating or, at the organismal level, blocks development. Identification of essential proteins and their characterization has yielded important insights into the functions of different cellular systems (2–5). However, the indispensability of a certain protein is a complex phenomenon, which may have several causes. At the cellular level, the lack of some essential proteins may create insurmountable problems for cell division, while deficiency of others may trigger cell death, i.e. spontaneous or programmed catastrophic failure of the cellular machinery. Importantly, cell death is most likely an integrative process that is realized via the complex interactions between different cellular systems (6). So, systematic understanding of triggers of cell death at the level of individual genes as well as gene interaction networks that mediate death can be very informative for a systems-level understanding of the cell.

Cell death is known to proceed via distinct mechanisms. Specifically, studies performed with mammalian cells have revealed more than 30 types of cell death (7), although they can be grouped into several larger categories (8). Even though many of the mechanisms involved in cell death in response to various pathologies and conditions have been established, specific and systematic understanding of the genes and systems whose perturbation can trigger necrotic death remains poorly understood. A recent systematic review on the subject of external treatments that can cause different types of cell death in yeast and other fungi (9) highlights that while numerous acute treatments are known to stop cell division and kill cells, little is known about whether and how cells die in conditions where division is not arrested.

The budding yeast *Saccharomyces cerevisiae* has been an immensely useful model in the analysis of lifespan and aging (10, 11), and also is an established model for the analysis of cell death (12–15). However, even in this simple and tractable model, there have been no systematic studies on genes whose perturbations can increase death rate. Consequently, there is also no understanding of the temporal dynamics of cell death during chronic or acute perturbation of different genes.

High-throughput testing of mutants for various phenotypes in microorganisms is very efficient, due to the ability of cells to grow in colonies on solid medium, which can be arranged in ordered and dense arrays on single plates. Colonies of yeast and other microbes are dense agglomerations of cells, which might seem like simple lumps of cells, but they are actually quite complex structures. This is especially true for wild yeast isolates, but even the laboratory strains, which lose the ability to form colonies with visible morphological complexity (16), exhibit various aspects of cell differentiation when grown in a colony (17). A probable reason for this diversity is that the colonies create gradients of various conditions from the outside surface to the internal regions, which, in turn, affect the behavior of cells in each region. However, this diversity has not been probed in a systematic fashion, and only fragmentary data are currently available in yeast (17–19), while studies of this phenomenon in prokaryotes have been more active (20). Currently, there is almost no data on how the location of yeast cells in colonies may affect their susceptibility to various perturbations, although this may be highly relevant for the killing of various pathogenic fungi using antifungal drugs.

In this study, we identified genes, whose perturbation increases the rate of cell death accompanied by membrane permeabilization, i.e. necrotic cell death, in dividing cultures by using the nontoxic stain phloxine B (21–23). The identified mutants exhibited distinct spatial patterns of death in colonial multicellular structures. We further developed novel tools to characterize the temporal dynamics of cell death in he identified mutants. These methods allowed us to determine that, in many mutants exhibiting increased death rates, mother and daughter cells may rapidly die in a stochastic manner during budding and cytokinesis and that this type of death seems to be related to perturbations in the functioning of the cell wall and membrane. Surprisingly, this type of death could be mitigated by neutral pH stabilization of the medium. Our discovery of this novel mode of necrotic death showed that cell division is a highly sensitive period of development and suggests that this type of death may be a common consequence of the dysfunction of various genes and the respective proteins, making this finding highly relevant to our understanding of various pathological states.

## Results

### Phloxine B staining of yeast colonies as a high-throughput method for detection of cell death

In order to identify mutants with increased rates of cell death, we employed the yeast knock-out (24) and essential gene knock-down (DAMP) (25) collections. To identify colonies with increased numbers of dead cells we adapted the use of phloxine B (Figure 1A) (21, 22) for high-throughput screening. Phloxine B is a negatively charged dye that only enters cells with impaired membrane permeability, i.e. those that have experienced necrotic death. Thus, if a colony contains mostly live cells, phloxine B would not stain this colony. However, if a colony has live cells (all colonies have live cells, otherwise they would not grow) as well as a substantial number of dead cells, these dead cells (Figure 1B) and therefore the colony would be stained in various shades of red. Plates containing phloxine B with yeast mutants were imaged after 2 days of growth (Supplemental archive 1), and the mutants exhibiting clear phloxine B staining were collected and replated onto new phloxine B plates, along with control strains, confirming the phenotype (Figure S1). To avoid population heterogeneity due to the emergence of suppressor mutants, all selected strains were streaked to single colonies with identical phenotypes, and further work was done on the progeny of single phloxine B positive colonies taken periodically from cryostorage. Despite this, while working with these phloxine-positive mutants, periodically strains stopped growing due to unknown reasons, thus in some of the tests presented further, the number of tested strains differs.

**Figure 1.**
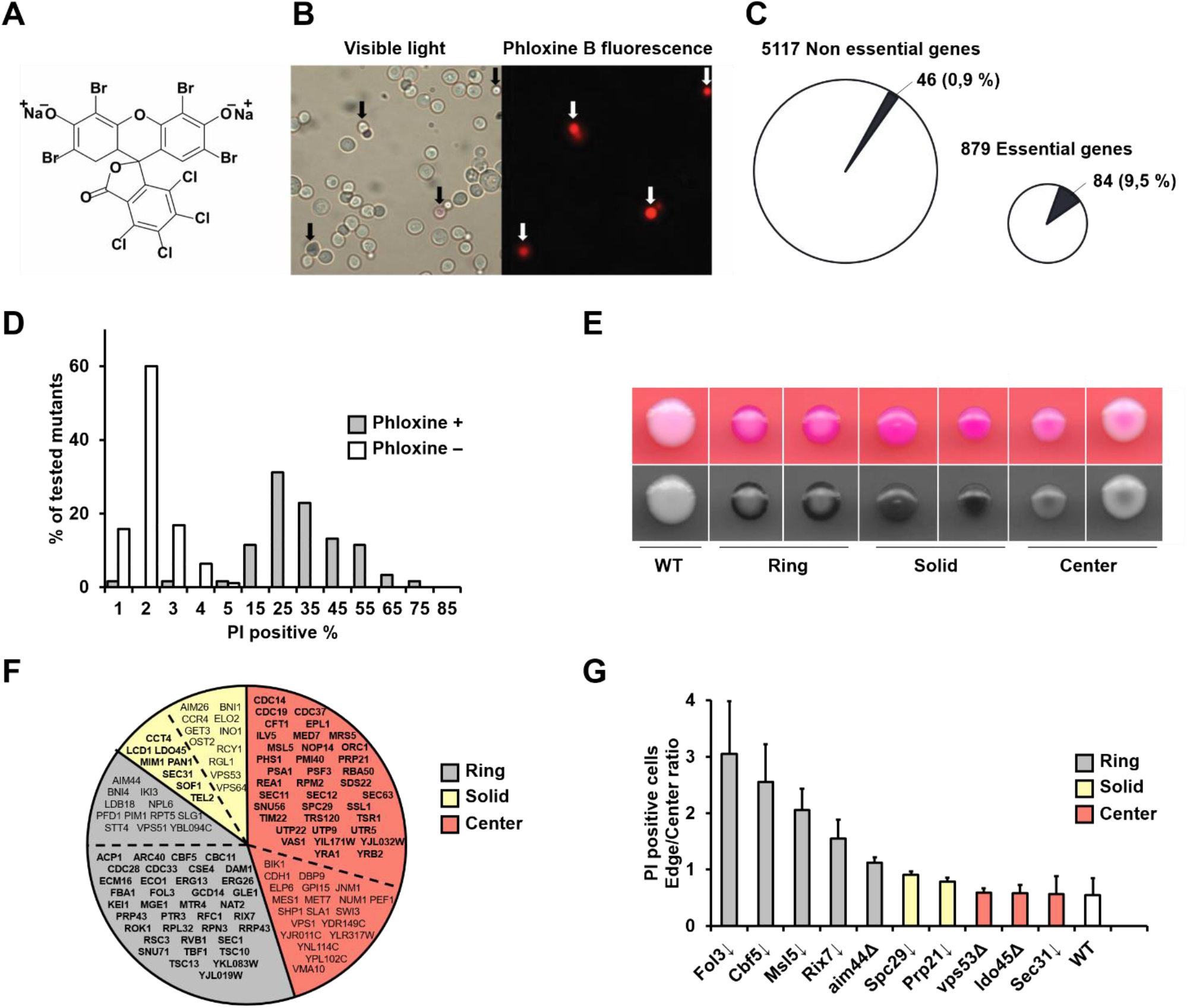
Phloxine B staining of mutant colonies provides spatial information on the distribution of dead cells. (A) Structure of phloxine B. (B) Fluorescent microscopy of live and dead cells stained with phloxine B. Arrows indicate dead cells. (C) Numbers of phloxine B positive mutants among the mutants with deletions of non-essential genes and downregulation of essential genes. (D) Distribution of the percentage of PI positive cells, obtained by a flow cytometric analysis of suspensions of yeast cells stained with PI. The number of phloxine B positive mutants was 61 and phloxine B negative mutants and wild type strain was 95. Measurements were performed in stationary cultures at least 2 times for each strain. (E) Colonies of the representative staining types grown for 48 hours on YPD containing phloxine B, depicted under the colony images. Specific mutants are (from left to right) – wild type, and downregulation of *CDC33, CDC11, SPC29, UTR5, SEC31*, and *RSC58* genes. Top panel – natural color, lower panel – green channel which provides maximal contrast for viewing phloxine B staining. (F) Genes associated with increased phloxine B staining. Colored sectors depict the phenotypic class (Ring, Solid, Center), regular font shows non-essential genes, and essential genes are shown in bold. (G) Ratio of the percentage of PI positive cells in cell suspensions obtained from the edge and center of yeast colonies grown for 48 hours on YPD medium. 3 colonies were analyzed for each strain, and error bars depict standard deviation.

Overall, we identified a total of 126 mutants in both collections, with 83 originating from the collection of mutants with downregulated essential proteins (out of 879 strains), and 43 from the knockout collection (out of 5,117 strains) (Figure 1C, Dataset S1, Figure S1). Thus, essential genes were strongly overrepresented among the genes whose dysfunction increases the chance of cell death.

To verify that phloxine B staining provides data comparable to the more commonly used methods of detecting dead cells, we used propidium iodide (PI) staining, which stains nucleic acids in cells with permeable membranes. For this, we tested 61 of our phloxine B positive mutants, as well as a panel of 90 random phloxine B negative mutants from the DAMP collection. We found that phloxine B negative mutants exhibited fewer than 4% PI positive cells, whereas more than 80% of the tested phloxine B positive strains contained more than 15% PI positive cells (Figure 1D). Thus, the use of phloxine B staining of colonies offers a qualitative, high-throughput assay that provides results in agreement with PI staining of *S. cerevisiae* liquid cultures.

We considered the possibility that increased staining could be related to an altered stain uptake or its removal by the yeast cells rather than due to cell death. To check this possibility, we tested whether phloxine B staining was dependent on the action of drug efflux pumps. Comparison of a wild-type strain and a mutant with deleted *PDR3* and *PDR5* genes, which are the main efflux pumps in yeast, showed that the latter exhibited slightly reduced, rather than increased, phloxine B staining compared to wild type (Figure S2).

### Phloxine B reveals spatial patterns of cell death in yeast colonies

One observation that immediately attracted our attention during the screen was that phloxine B staining patterns differed between the colonies of different mutants (Figure1E, Figure S1). A large number of strains exhibited a pattern of staining where only the border (edge) of the colony was stained, while the center was much lighter (hereafter called “Ring” mutants). Other mutants exhibited more or less homogenous staining (“Solid” mutants), while the third class of mutants exhibited staining mostly in the center of the colony (“Center” mutants), albeit mutants with a strong phenotype like this were rare. Overall, out of the 126 mutants, 51 possessed the Ring phenotype, 56 Solid, and 19 Center phenotype (Figure 1F, Dataset S1).

In order to exclude phloxine B-specific effects, such as toxicity or diffusion gradients, colonies were grown on medium without phloxine B, after which cells were collected from the edge and center of the colonies and assayed for the percentage of cells stainable by PI. The ratio between these percentages represents the degree to which more dead cells are present on the outside of the colony as opposed to the center and were in excess of 1.5 for the Ring mutants, around 0.5 for the Center mutants, and close to 1 for the Solid mutants (Figure 1G). Colonies of the wild type strain had very few dead cells, and thus the results were highly variable; however, the ratio suggested that in wild type colonies, dead cells might be more numerous in the center. Overall, the data revealed that phloxine B staining offered a convenient qualitative test for determining the spatial distribution of dead cells in yeast colonies.

### Functional analysis of genes whose perturbation increases cell death

Using a GO-term enrichment analysis of genes whose perturbation increases cell death, we observed enrichment of genes associated with cell division (including those involved in the cell cycle, as well as tubulin and actin cytoskeleton) and establishment of [component] localization in the cell (Figure 2A). The cell cycle and cytoskeletal categories mostly overlapped; however, the localization category overlapped with the latter two only partially (Figure 2A). Genes belonging to these categories accounted for ∼45% of the genes found to increase cell death.

**Figure 2.**
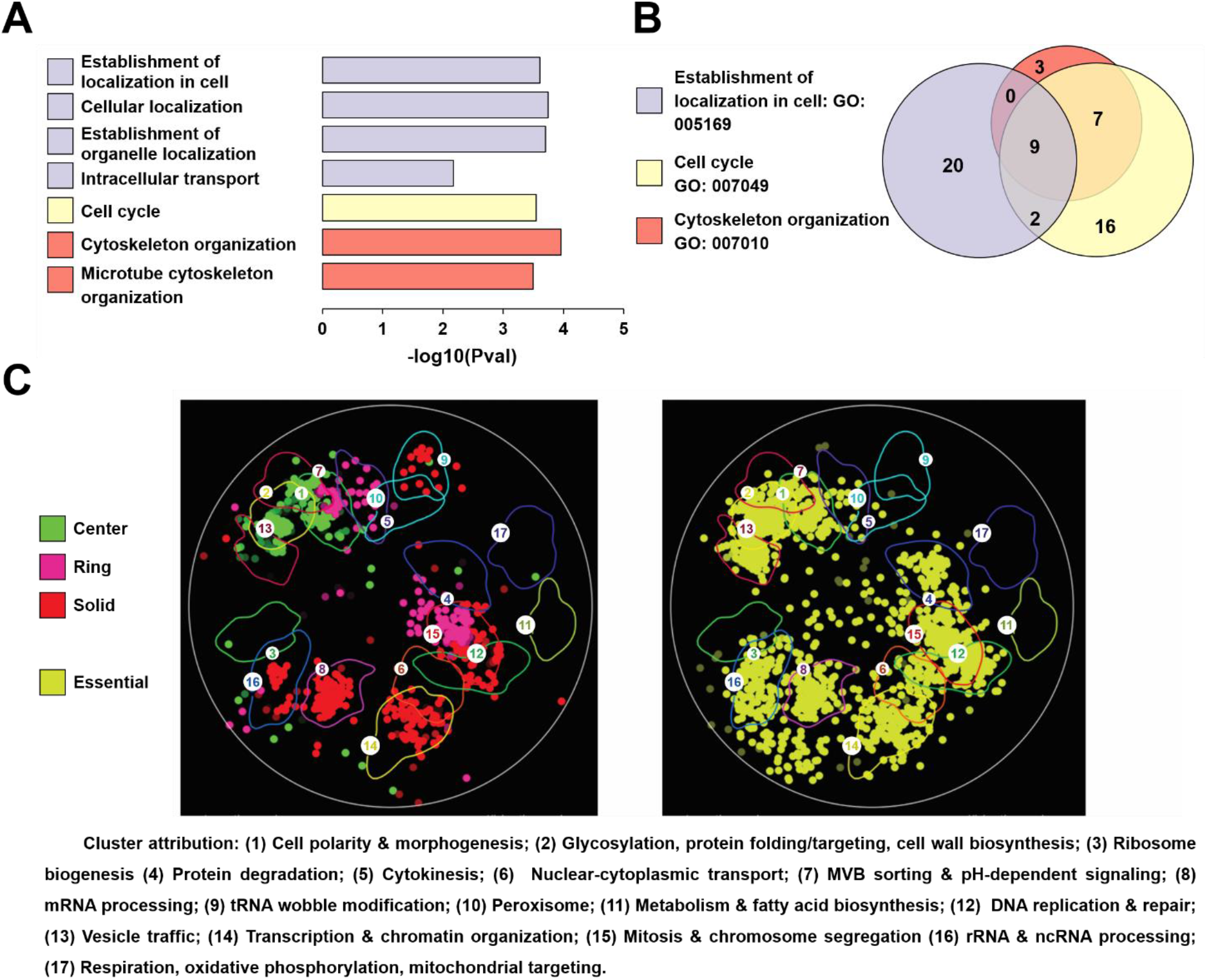
Enrichment analysis of genes identified by phloxine B staining. (A) GO-term enrichment analysis. Bars with the same color denote GO-terms which are nested into each other. (B) Venn diagram of the identified GO-term enrichment with the number of genes in each sector. (C) SAFE-analysis of the identified genes presenting specific phloxine B staining patterns with color coding (green – center, magenta – ring, red – solid) (left) and all essential genes from the DAMP collection (yellow) (right). Spots indicate genes that represent nodes of the interaction network enriched for interactions with the input list of genes. Spot brightness indicates p-value of enrichment.

We further employed the SAFE algorithm (26), which is a method of identifying functional enrichment of gene sets based on the similarity of experimentally mapped genetic interaction networks (27) rather than manually assigned gene annotations. SAFE revealed enrichment in interactions with genes involved in DNA replication & repair, mitosis & chromosome segregation, cell polarity and morphogenesis, transcription & chromatin organization categories, which agrees with the GO term analysis results, as well as other categories. Notably, enrichment patterns for different phloxine B staining types were distinct, with the Center phenotype forming a compact cluster in the regions corresponding to vesicle traffic, glycosylation & protein folding/targeting & cell wall biosynthesis; MVB sorting/pH-dependent signaling; cell polarity and morphogenesis, while the genes whose perturbation causes the Ring phenotype were mostly found in the regions corresponding to MVB sorting/pH-dependent signaling and mitosis & chromosome segregation. For the Solid phenotype, interacting genes clustered with a wider range of categories - tRNA wobble modification; mRNA processing; rRNA and ncRNA processing; mitosis & chromosome segregation, DNA replication & repair and transcription & chromatin organization. Overall, the SAFE distribution for mutants identified by phloxine B screening was similar to that of essential genes from the DAMP collection (Figure 2C, right panel), even though 1/3 of our dataset consisted of non-essential genes (Figure 2C, left panel). Notable differences included the absence of enrichment for the protein degradation category among the mutants exhibiting phloxine B staining, and the emergence of the tRNA wobble modification category, which was absent in the essential gene enrichment pattern.

### Pervasive death of newborn cells

It is unclear whether the dead cells accumulating in colonies and liquid cultures resulted from cells dying quickly after birth, or if cell death occurred after a prolonged period of division arrest. In order to answer this question, we devised an assay to detect death of recently born daughter cells (Figure 3A). In brief, cells were labeled with a membrane-impermeant fluorophore that stains amino-groups in the cell wall, and the cells were allowed to divide 1-2 times. This generated a population of unstained replicatively young cells, whose chance of death can be determined by a flow-cytometric analysis of PI staining (Figure 3B). We termed this method the DON assay (Death of Newborns). DON supported a clear distinction between mutant and wild type strains and demonstrated that a notable percentage of cells born experienced death during the first 1-2 divisions, wherein membrane permeabilization occurred relatively quickly after appearance of these cells. We found that for ∼95% of our identified and tested mutants (n=77), young mutant cells were at least 3-fold more likely to die compared to the situation in wild type cells (Figure 3C, Dataset S2). Aging cells also seemed to experience death; however, interpreting the relative chance of cell death between young cells and aged cells is difficult, because a portion of the dead stained cells originate from cells that were dead during staining, i.e. the results are not easily interpretable without more time points. This study is currently underway and will be published at a later time.

**Figure 3.**
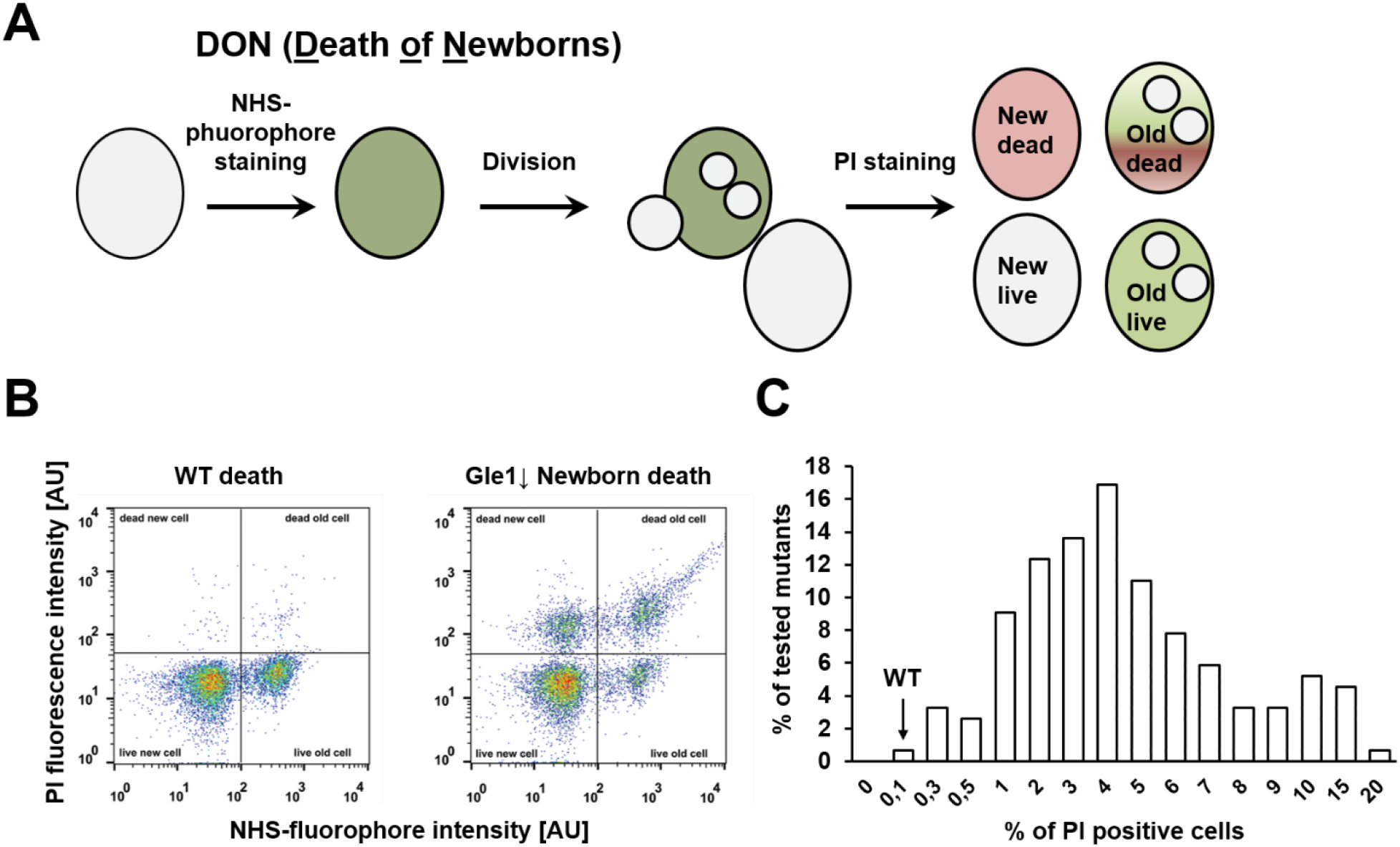
Phloxine B positive mutants exhibit increased chance of rapid death in early life. (A) Schematic of the method to assay death of newborn cells (DON). (B) Scatter plots of cells not exhibiting (left) or exhibiting (right) considerable death of newborn cells, as obtained by flow cytometry. (C) Distribution of percentage of PI-positive cells in various tested mutants and WT strain in young cells as assayed using the DON method (number of analyzed mutants – 77). Each strain was analyzed twice in independent experiments, with a similar trend being observed.

### Real time analysis of cell death

In order to monitor the death of both mother and daughter cells, as well as to determine how quickly cell death occurs and whether there are some changes to the cell prior to death, we employed real-time microscopy in the presence of phloxine B. We also tested whether PI/phloxine B staining might occur transiently, without cell death, as reported for PI staining under specific circumstances (28–30). Thus, we monitored the division and permeabilization of cells in the presence of phloxine B. Due to the labor intensity of microfluidic assays, we analyzed only 5 strains: 4 phloxine B positive strains, and a WT strain.

First, we found that strains selected based on colony staining indeed commonly exhibited cells that acquired phloxine B staining during culturing (Figure 4A and Videos 1 and 2). We did not observe any cells that proceeded through division after they had acquired robust phloxine B staining. We also found that phloxine B acquisition correlated with increased transparency in the phase contrast channel (Figure 4C, D), indicating a considerable change of phase properties, as was noted previously during the quantitative phase contrast microscopy study of yeast death in response to plant defensin treatment (31). The same is also observed in human cells (32). This increased transparency was somewhat reduced after it appeared, while the phloxine B signal increased and persisted indefinitely. Notably, cells stained with phloxine B did not disappear or experience complete lysis in any of the tested strains during our observations. Also, we did not observe any phloxine B negative cells that arrested division, which suggests that no other, non-permeabilizing type of death or probabilistic shift into quiescence took place in the studied strains (Dataset S3).

**Figure 4.**
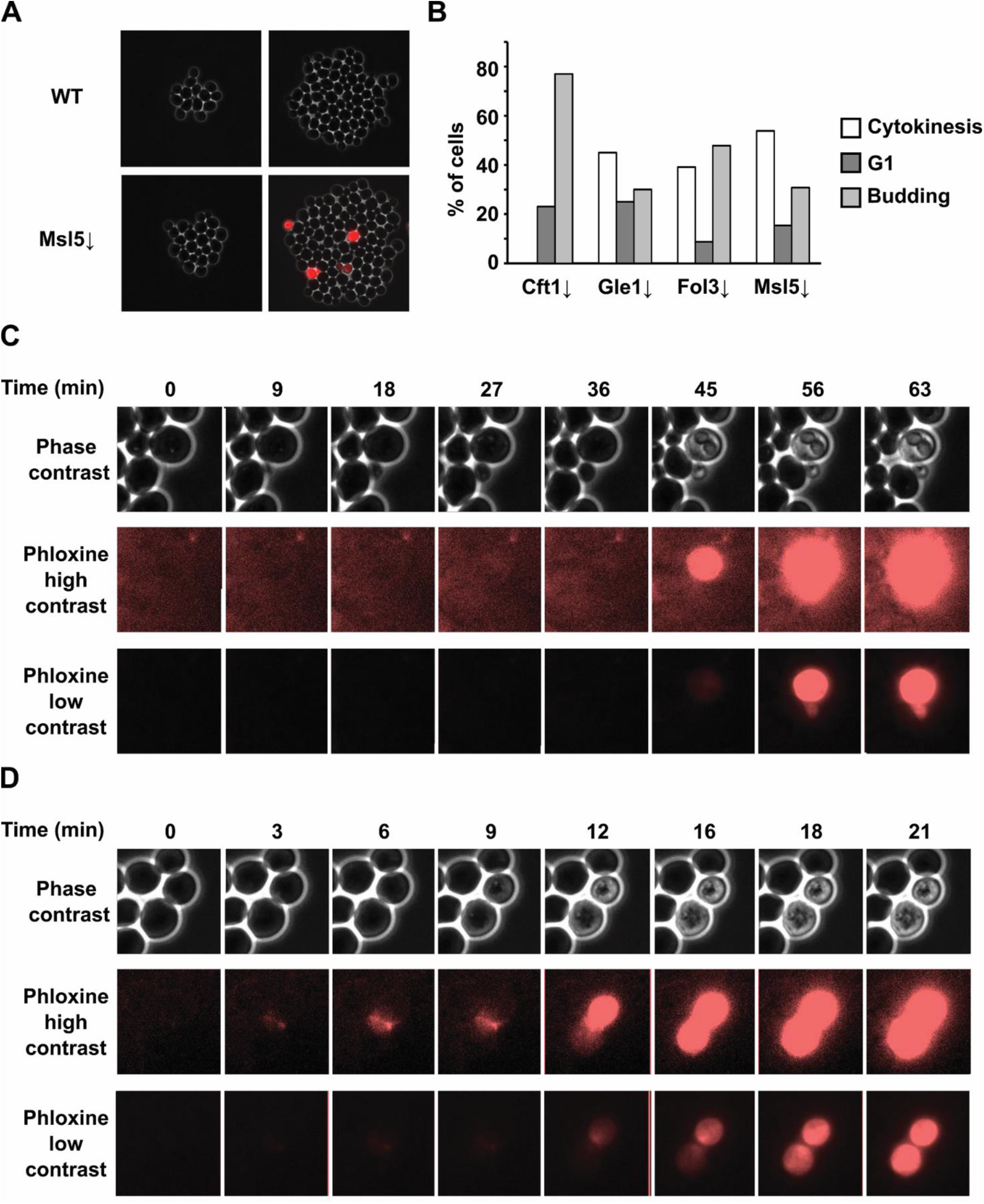
Real-time visualization of cell death accompanied by membrane permeabilization. Cells were grown in a microfluidics chamber supplied with YPD medium containing phloxine B and visualized using differential interference contrast and fluorescence imaging. (A) Microcolonies of yeast at two different timepoints. (B) Histogram of the approximate cell cycle stages of cell death for different mutants (number of analyzed dead cells – CFT1 – 24; GLE1-27; FOL3 – 34; MSL5 - 32. (C) Time lapse images of a mother dying with a small bud for a mutant with *CFT1* knockdown (See video 1). (D) Time-lapse images of a mother-daughter pair of cells which die during cytokinesis for the mutant with *MSL5* knockdown (See video 2 and video 3). All time lapse experiments were performed in two independent runs for each strain, with no less than 4 chambers, i.e. microcolonies, analyzed for each strain during a run. A control strain was also included into each run.

A common occurrence in the phloxine B positive strains was that they exhibited high levels of cell death at the budding stage. For all of the tested strains, this represented 30-75% of the dead cells observed (Figure 4B, C and Video 1). Also, cells quite often died shortly after cytokinesis (Figure 4B, D and Video 2). Notably, in both cases, death could sometimes be observed only in the daughter cell, only in the mother cell, or in both the cells simultaneously without any discernible pattern.

The data indicate that budding and cytokinesis are high periods of the yeast lifecycle, when perturbation of different systems seemingly unrelated with these processes can increase the likelihood of a catastrophic failure.

### Phloxine phenotypes are strongly associated with indications of membrane and cell wall dysfunction

Why would bud generation and cytokinesis be vulnerable stages of the life cycle? We hypothesized that during these stages, the budding yeast cell-pair (mother and daughter) needs to rapidly and precisely expand and/or remodel both the plasma membrane and the cell wall. In order to test whether impairments in these organelles might be common among phloxine B positive mutants, we tested 76 of these mutants, as well as a control subset of phloxine B negative mutants from the DAMP collection for sensitivity to growth on SDS and ethanol (associated with effects on both the plasma membrane and lipid metabolism) (33–35) as well as to two cell wall stressors: Calcofluor white (CFW), which interacts with chitin (36) and other cell wall carbohydrates, and Congo Red, which interacts with β-1,3-d-glucan (37). We observed that for a set of phloxine B positive mutants with reduced abundance of essential proteins, ∼25, 45 and 25% of the strains were sensitive to SDS, ethanol or CFW, respectively (Figure 5 and Figure S3, Datset S4), while almost none from a randomly selected set of 90 phloxine B negative mutants from the same collection exhibited sensitivity. Notably, Congo Red showed no dramatic difference between the number of sensitive mutants between the phloxine B positive and negative sets (Figure 5). These observations suggest that phloxine positive mutants are specifically sensitive to some stressors targeting the cell membrane and cell wall, but not to all of the tested stressors.

**Figure 5.**
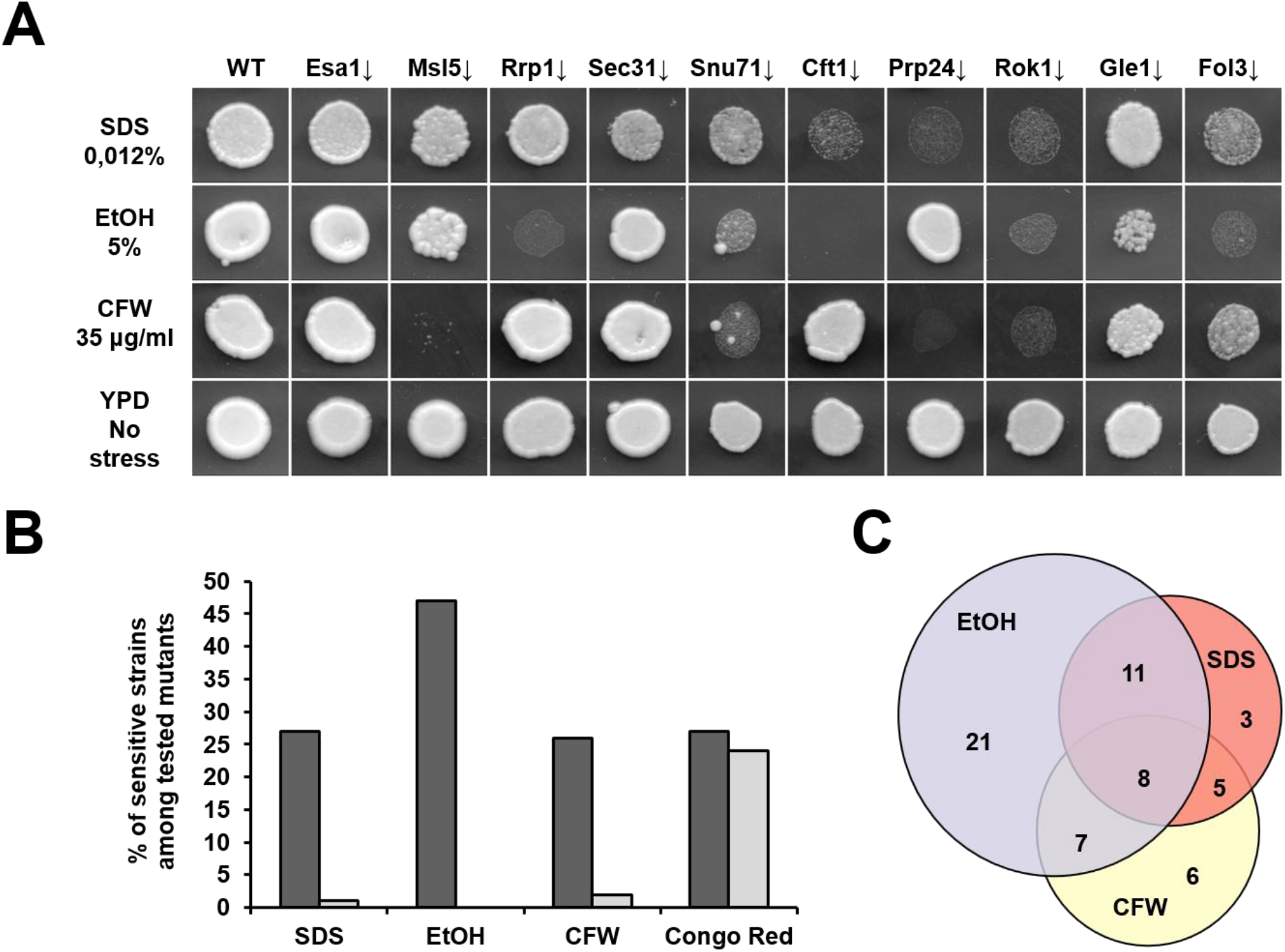
Phloxine B positive mutants with reduced abundance of essential proteins are sensitive to membrane/cell wall stress. (A) Determination of stress sensitivity of indicated Phloxine B positive mutants. Complete data are available in Figure S3 (76 mutants tested) (B) Phloxine B positive mutants (dark shading) show high incidence of sensitivity to ethanol, SDS, Calcofluor white, but not Congo Red respectively, as compared to a control set of non-phloxine staining mutants from the same mutant collection (90 mutants tested) (light shading). (C) Venn diagram of the stressor assays with the number of sensitive mutants in each sector.

### Identification of universal features of phloxine B positive mutants and mitigation of death caused by different mutations

A wide range of literature on cell death in response to various stimuli in yeast is available and a common feature of death is the generation and causative role of reactive oxygen species (ROS) in death. Lipid peroxidation in the membrane might also explain the commonality of membrane impairment in various mutants. Thus, to see whether reactive oxygen species (ROS) played a role in the cell death observed in our study, we used the ROS-sensitive stains dihydroethidium (DHE) (89 mutants tested) and 2’,7’-Dichlorodihydrofluorescein diacetate (DCFDA) (79 mutants tested) to detect presence of oxidative stress among phloxine-positive mutants. This revealed increased levels of ROS (>2-fold) in only 11 of the tested mutants for DHE staining, and 9 for DCFDA) (Figure 6A and Datasets S5 and S6). However, these mutants did not show a lowered chance of death in response to the supplementation with N-acetyl-cysteine, rather they were often more sensitive, sometimes quite dramatically so (Figure 6B). In order to get a more complete understanding of the effect of this antioxidant on cell death in phloxine-positive mutants, we tested 79 mutants, and observed that only 1 of them (Mdn1↓) exhibit a more than 2-fold reduced cell death in response to medium supplementation with NAC (Dataset S6). Notably, this strain did not exhibit increased staining with DHE or DCFDA. This suggests that in most of the tested phloxine-positive mutants, death is not caused by oxidative stress, while in the few strains that do exhibit increased ROS levels, this increase is not a pro-death factor, but might actually be an adaptive response.

**Figure 6.**
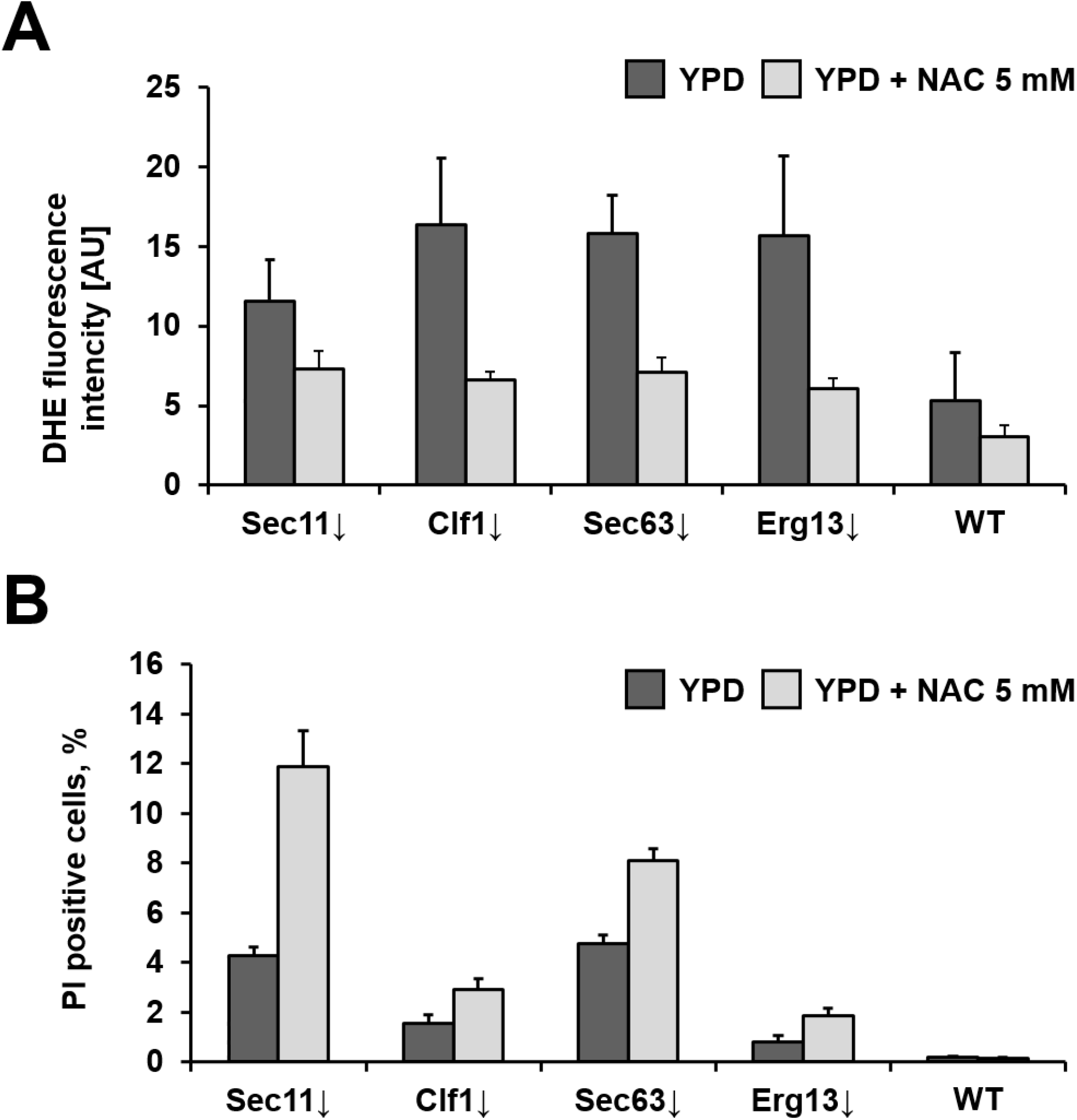
Cell death of phloxine B positive mutants with chronic oxidative stress is not prevented by N-acetyl cysteine (NAC). (A) The presented Phloxine B positive mutants exhibit increased staining by DHE in living (Sytox-green negative) cells (identified by screening 89 mutants, see figure S4). This staining is mitigated by N-acetyl-cysteine (NAC) treatment. (B) Treatment with NAC does not reduce, but rather stimulates cell death in the strains exhibiting increased DHE fluorescence in living (PI-negative) cells.

After that, we tested a range of other treatments that might have mitigating effects on cell death. This was done by plating 81 identified phloxine B positive mutants onto YPD with phloxine B supplemented by various compounds (Figure 7A). Some of the tested treatments indeed mitigated cell death but only in a limited number of mutants and this data might be useful for a deeper understanding of cell death mechanisms in future studies (Figure S5). However, two treatments had a strong mitigating effect on nearly all of the mutants – these were the use of potassium acetate as a carbon source, and buffering the medium to pH 7. Because cultivation in potassium acetate increased external pH and other non-fermentable carbon sources (ethanol, glycerol) did not mitigate the phloxine B phenotype (data not shown), we concluded that the effect of potassium acetate and pH buffering were one and the same, and were due to the neutral external pH. Unlike all of the other tested treatments, whose effects were limited to ∼30% of the mutants or less, increased external pH reduced the phloxine B phenotype in most of the tested mutant colonies (Figure 7A, Figure S5, Dataset S7). Because of the possibility that phloxine staining is pH-dependent irrespective of cell death rates, we also verified our results by testing cells grown at buffered pH without phloxine B, but then washed the cells (to control for the effect of external pH on the staining) and stained with PI (Figure 7B), confirming the results obtained using phloxine B.

**Figure 7.**
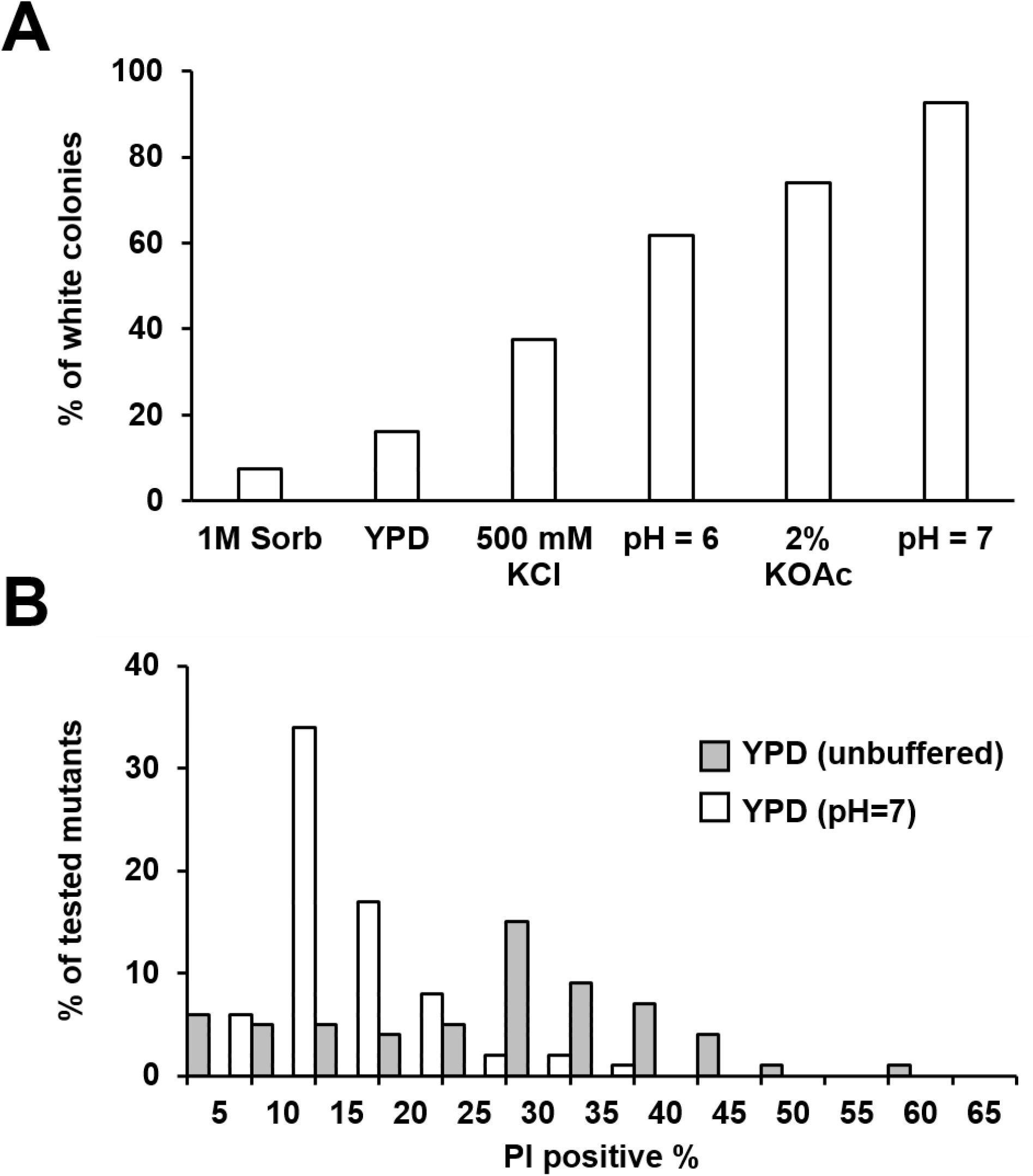
Various treatments reduce phloxine staining in specific mutants, with external pH stabilization having a near-universal effect. (A) 81 mutants exhibiting varying degrees of phloxine staining were plated onto YPD medium containing phloxine B supplemented with the denoted component (except for 2% KOAc, in which 2% potassium acetate was used as a carbon source instead of glucose). The graph denotes the share of tested mutants with a non-phloxine B staining phenotype. Sorb – sorbitol, KCl – potassium chloride, pH=6 – buffering with 20 mM phosphate buffer; KOAc – potassium acetate as the sole carbon source; pH=7 buffering with 20 mM phosphate buffer (B) The same mutants (see (A) and Figure S5) were grown in normal YPD or YPD buffered to pH=7. They were subsequently stained with PI and the % of PI-staining cells was measured by flow cytometry.

Thus, our results show that neutral external pH has a near-universal mitigating effect on the occurrence of necrotic death associated with division. We hypothesized that breaching of the cell membrane might foremostly be deadly due to cytoplasmic acidification. If this was case, then at lower pH, membrane permeabilization should be irreversible, as observed above, while at higher pH, where no acidification occurs during rupture, the membrane damage could be repaired, and manifest as a transient permeabilization event. Using microfluidics, we tested this hypothesis but did not detect any transient permeabilization events (data not shown).

## Discussion

In this study, we performed the first systematic identification of genes, both essential and non-essential, whose perturbation increases the rate of necrotic cell death involving membrane permeabilization in dividing *S. cerevisiae*. While the overall populations of identified mutants exhibited an increased chance of cell death, most cells were capable of multiple divisions. We considered a possibility that, for downregulated essential genes, this stochastic effect may be caused by the unequal downregulation of gene expression and therefore protein levels in different cells of the colony; however, the same effects were seen for the gene deletion mutants, where all cells are equally targeted. Thus, while stochastic cell death might be influenced by the heterogenous expression of target genes, it should also be related to some other forms of cell to cell variation or probabilistic events.

Our results clearly demonstrate that perturbation of numerous cellular systems may result in an increased chance of necrotic cell death. The identified genes and their enrichment patterns closely followed the enrichment exhibited by essential genes (Figure 2C), even though a third of the genes identified in this work were non-essential. However, some gene categories that represent a sizeable fraction of essential genes were absent among the mutants we identified. For example, according to the SAFE analysis, essential genes are enriched in the protein turnover category, which was not detected in our data (Figure 2C). Apparently, impairment of genes in this category does not result in necrotic death or alternatively this impairment needs to be more severe than what could be achieved by this type of downregulation (25).

Since our screen revealed the overall enrichment of genes implicated in cell cycle, cytoskeletal functions, and organelle localization (Figure 2A), and we observed death during the small bud stage and shortly after cytokinesis (Figure 5 and Movies 1 and 2), we hypothesize that perturbations in various unrelated processes can affect the ability of the cell to repair or generate the cell wall or plasma membrane during bud formation and/or cytokinesis. Although this was not detected as a statistically significant enrichment, 10 out of the identified 126 genes are involved in lipid metabolism, which is a key process for membrane synthesis and remodeling, with 6 more were reported to be sensitive to ethanol (38), a phenotype associated with impaired plasma membrane (39). Notably, changes in lipid synthesis have been reported to occur during the cell cycle in yeast (40) and mammalian cells (41). We were also able to obtain additional support for the impairment of the membrane and cell wall by characterizing specific sensitivities to membrane- and cell-wall disrupting compounds. On the other hand, sorbitol, which often reduces toxicity of cell-wall perturbing compounds, did not show a mitigating effect on cell death in most mutants. This finding suggests that overall breakdown of the cell wall is unlikely to be the direct cause of cell death. However, the commonality of CFW sensitivity among phloxine B mutants suggests that some functions that involve chitin might be involved. This is also highlighted by the observations that stochastic necrosis is associated with cytokinesis, which involves deposition of reparative chitin at the site of bud/birth scar formation.

Our ability to differentiate between spatial patterns of cell death in colonies allowed us to uncover differences between cell death caused by distinct genetic perturbations. Mutants prone to death on the colony edges exhibited enrichment of genes belonging to the mitosis and cell polarity/morphogenesis categories. This suggests that the cells located on the bottom and edge of the colony, and thus experiencing the highest availability of nutrients and exhibiting rapid division (18, 42) are more prone to death in response to the impairment of genes involved in cell division.

Mutants prone to homogenous death throughout the colony exhibited enrichment in genes belonging to the tRNA wobble modification cluster, which was not observed for other groups of phloxine B positive mutants and may reflect regulation of cell proliferation based on tRNA abundance and mRNA codon usage (43, 44). Interestingly, mutants exhibiting this pattern of dead cell distribution were also enriched in the genes at the intersection of mitosis and DNA-repair and replication clusters. This suggests that some aspects of DNA repair might be important for survival of both rapidly dividing cells on the outside of the colony, as well as slowly dividing or non-dividing cells in the colony center.

Mutants prone to death at the center of a colony showed enrichment in genes related to vesicle transport and glycosylation/protein folding/cell wall categories, suggesting that damage to these machineries is especially relevant to starving and/or respiring cells known to inhabit the deeper regions of colonies (42). Interestingly, this pattern of cell death has previously been reported in yeast colonies growing on non-fermentable carbon sources for prolonged periods of time (17).

Overall, different patterns of cell death in the colonies demonstrate that dividing and non-dividing cells, or cells with different physiology might have very distinct sensitivities to diverse death stimuli, i.e. it might be possible that cells at the colony center might be physically or chemically shielded by cells in direct contact with the medium. Such phenomena have been reported previously, including nutrient sharing between dying cells on the inside of a colony and live cells on the edge (17) and the protective role of dead cells against the effects of polyene antifungal drugs (45).

Identification of mutants with an increased probability of cell death provided us with an opportunity to study dynamics of this novel type of cell death. One notable observation was that the death was most often rapid, as evidenced by the DON assay and microfluidics data (Figures 3 and 5 and videos). This is in contrast to the observation of a relatively slow onset of necrosis in senescent cells obtained during replicative aging (46, 47). Also, this is unlike the case of hydrogen peroxide stress, which, at lower concentrations, induces cell death via processes that do not involve rapid membrane permeabilization, and causes necrosis at higher concentrations (48). Because we did not observe any abnormal cellular features or division arrest prior to cell death, we assume that the death type is most likely primary necrosis occurring as a consequence of stochastic damaging events. This conclusion challenges the common view of necrosis being mainly a feature of severe stress, as opposed to apoptotic death, which occurs during milder perturbations.

To sum up, our work is the first to identify genetic perturbations that increase the chance of cell death in proliferating cells and, most likely, many of these mutations cause cell death associated with the process of cell division. They also demonstrate that seemingly functionally unrelated mutations cause cell death that can be mitigated, via an unknown mechanism, by stabilizing external pH near neutral level, and that mutants with this type of cell death often exhibit phenotypes associated with perturbed properties of the plasma membrane and/or cell wall. This suggests that different impairments to the complex architecture of the cell seem to have a universal downstream effect on some stochastically-dangerous processes, in this case –possibly the maintenance of optimal plasma membrane or cell wall condition during division. We also provide a set of new methods to study cell death: high-throughput screening and microfluidics based on phloxine B staining, which can detect increased rates of cell death under various conditions and distinguish different types of dead cell distribution in colonies, as well as the DON assay to detect rapid necrosis in young cells. Finally, our work identifies a large set of mutants that exhibit stochastic necrosis and may be characterized in greater detail in future studies, thereby leading to a better understanding of specific differences in the modes of cell death caused by particular genetic perturbations.

## Materials and Methods

### Yeast strains

Strains used in this work were derivatives of the BY4741 strain (*MATa his3Δ1 leu2-D0 met15 Δ0 ura3 Δ0*), obtained in the studies by (2, 25).

### Screening of genome-wide mutant collections

Strains from the tested collection were refreshed on YPD medium and then pin-spotted onto YPD with phloxine B (30 μM) (Acros Organics, Cat # 189470250) using 384-pin replicators with long pins (Singer, UK). The colonies were grown for 48 hours and then scanned using an Epson Perfection V550 Photo flatbed scanner at 2000 dpi. For optimal viewing of phloxine B staining phenotype, FIJI (49) was used to extract green channel images from the raw RGB, thus providing the highest contrast in terms of phloxine staining. Each library plate was tested at least twice and strains that did not show reproducible results were not taken up for further analysis.

### Gene enrichment analysis and SAFE

Gene enrichment analysis was performed using The GO Term Finder (Version 0.86) which can be found at https://www.yeastgenome.org/goTermFinder. Proportional Venn diagrams were constructed using BioVenn (50). SAFE analysis was done using tools described in (26) for separate lists of genes identified as having different phloxine B staining patterns, and then combined onto a single map.

### Flow cytometric methods

Flow cytometry was performed in 96-well plates using a Guava EasyCyte 8HT flow cytometry system (Millipore, USA) equipped with a 488 nm laser and a Cytoflex S equipped with a 405, 488 and 561 nm laser. Where applicable, cell fluorescence was measured in the green (525/30 nm) (for FAM fluorescence detection in the DON method and Sytox green fluorescence) and red (690/50 nm) channel (for PI and DHE fluorescence detection). For all experiments except those with ROS levels assayed by DHE, prior to analysis, cells were stained with PI (2 μg/ml) for 1 hour in distilled water. For detection of dead cells in experiments using DHE, Sytox Green was used in similar fashion.

### Determination of dead cell numbers in liquid cultures

Stationary liquid cultures were obtained by growing cells in 96-well plates with shaking at 30°C, stained with PI as noted above and analyzed using flow cytometry.

### Determination of differential death in yeast colony regions

Colonies of yeast grown on YPD medium for 48 hours were used for manual collection of cells from the outer edge and center of a colony using a sterilized wire loop. These cells were suspended in distilled water and then stained and analyzed as noted above.

### Death of Newborns Assay

Stationary cultures of yeast cells were grown in 96-well plates, then diluted by a factor of 50 with fresh YPD. After allowing sufficient time to divide 2-3 times, these cells were stained using FAM-NHS (Lumiprobe, Russia) to label the cell wall green. Cells from overnight cultures that were diluted 50-fold in fresh YPD and allowed to regrow for 5 hours were spun down in plates, washed twice with distilled H_2_0, spun down again and the supernatant was removed. FAM-NHS was added to each well (1 mM concentration) in a volume of 30 μl in 10 mM PBS (pH = 7.5). After 10 minutes of vigorous shaking in a plate covered with aluminum foil, the cells were washed 3 times with 120 μl of distilled water per each well. The obtained cells were then split into aliquots, one of which was kept for analysis as the zero time point, while the other was inoculated into YPD and allowed to accomplish a few divisions before flow cytometric analysis. The unlabeled cells were considered to be the youngest, because they were the ones that appeared after the staining procedure.

### Microfluidic-based real-time microscopy

Live-cell real-time microscopy was performed in a custom-made microfluidic device made of polydimethylsiloxane and a glass cover-slip that allows trapping of cells in a dedicated region of interest, limiting colony growth in the XY-plane. Constant medium flow at 20 µl/min was applied, enabling imaging of colony growth over several generations. Cells were cultured in YPD medium containing 300 nM Phloxine B. A Nikon Eclipse Ti-E with SPECTRA X light engine illumination and an Andor iXon Ultra 888 camera was used for epifluorescence microscopy. A plan-apo λ 100x/1.4Na Ph3 oil immersion objective was used to take phase-contrast and fluorescence images with a 3-minute frame rate. For automated focusing, the built-in Nikon perfect focus system was used during the experiment. Phloxine B fluorescence was imaged by exposure for 200 ms, illuminating with the SPECTRA X light engine at 556 nm and about 10 mW power. Identical settings were used for each of the experiments. Temperature control was achieved by setting both a custom made heatable insertion and an objective heater to 30°C.

### Measurements of ROS levels and determination of the effects of NAC

For testing of the levels of ROS using DHE and DCFDA, cells were grown overnight on solid YPD medium, after which they were resuspended in YPD (or YPD with 5mM NAC), incubated for 6 hours (with addition of Sytox Green or PI 30, as well as ROS-sensitive stain (final concentration - 10 μg/ml DCFDA; DHE – 5 μg/ml) 30 minutes prior to the end of the incubation), and then subjected to flow cytometry. ROS levels were only scored in cells negative for Sytox Green/PI.

For targeted testing of the effects of NAC on ROS levels and cell death (Figure 6), cells were grown to logarithmic phase in YPD or YPD with 5mM NAC, after which the levels of ROS and the amount of dead cells was measured using DHE and Sytox green together, and Propidium iodide alone, respectively.

### Detection of phloxine B staining under various conditions and testing of mutant sensitivity to various compounds

A subset of the identified phloxine positive mutants (Figure S4 and Dataset S6) as well as the control wild type strains were plated onto YPD medium containing phloxine B or the same medium supplemented with various compounds (1M Sorbitol, pH buffers). For sensitivity testing, we first determined the maximal concentration at which the wild-type strain showed robust growth on YPD medium supplemented with stressors (SDS, Ethanol, CFW, Congo Red), and then plated the subsets of phloxine positive and negative mutants (Figure S3 and Dataset S4) onto plates with these identified concentrations. Phloxine phenotype and growth sensitivity was scored manually. Each experiment was performed at least twice.

## Supporting information

Supplemental data, including a description of all the data and figures, datasets and videos

## Acknowledgements

S.E.D. is a member of the Interdisciplinary Scientific and Educational School of Moscow University “Molecular Technologies of the Living Systems and Synthetic Biology”.

## Funding

This project was partially funded by a grant from the President of the Russian Federation for Young Scientists MK-3323.2019.4 (initial screening of the deletion collections, DON method and microfluidics), the Russian Science Foundation grant #21-74-10115 (identifying conditions that prevent cell death, study of sensitivity of phloxine B mutants to stressful treatments, study of ROS-related effects) and the Ministry of Science and Higher Education of the Russian Federation (base funding).

## Author contributions

AIA conceived and performed the experiments, analyzed data, drafted the manuscript, contributed funding and equipment, EVG performed experiments, analyzed data and created figures, OVM performed experiments, AVN analyzed data and created figures, IVK performed experiments and aided in data analysis, RS provided equipment and analyzed data, ESS performed experiments and provided equipment, SED analyzed data, discussed the results, and edited the manuscript, VNG conceived the study, discussed the results, and edited the manuscript.

## Notes

### Competing Interest Statement

The authors have declared no competing interest.

### Summary of Updates

Corrected errors (figure numbering), added data (on the role of antioxidants in modulating cells death), small textual revisions

