## Supplemental data, including a description of all the data and figures, datasets and videos for "Systematic identification of yeast mutants with increased rates of cell death reveals rapid stochastic necrosis associated with cell division": Supplemental figures and materials.docx

**Supplementary Data**

**Dataset S1.** Tables of genes whose perturbation causes phloxine staining and maps of testing plates

Sheet 1 - List of genes, whose deletion/downregulation increases phloxine staining

Sheet 2 – Map of mutants in Plate I (phloxine positive mutants from DAMP collection)

Sheet 3 – Map of mutants in Plate II (phloxine positive mutants from DAMP and KO collection)

Sheet 4 - Map of mutants in phloxine negative mutant plate

Excel file

**Dataset S2. Data on DON assay**

Excel file

**Dataset S3. Dynamics of cell division obtained from microfluidic observation of single cell division.** Each sheet is the data obtained for a single microcolony of a microfluidics experiment on a strain with a noted downregulated gene. Almost no cells were observed to arrest division in the absence of phloxine B staining. Cells stained with phloxine B were not included.

Excel file

**Dataset S4. Tabulated data on the growth phenotypes of phloxine positive mutants on medium with different stressors.**

Excel file

**Dataset S5. Tabulated data on flow cytometric analysis of phloxine positive mutants for DHE staining efficiency, which was used to select strains with reliably increased staining.**

Excel file

**Dataset S6. Tabulated data on phloxine phenotypes of plate I mutants (DAMP mutants) on media with different, potentially mitigating, additives.**

Excel file

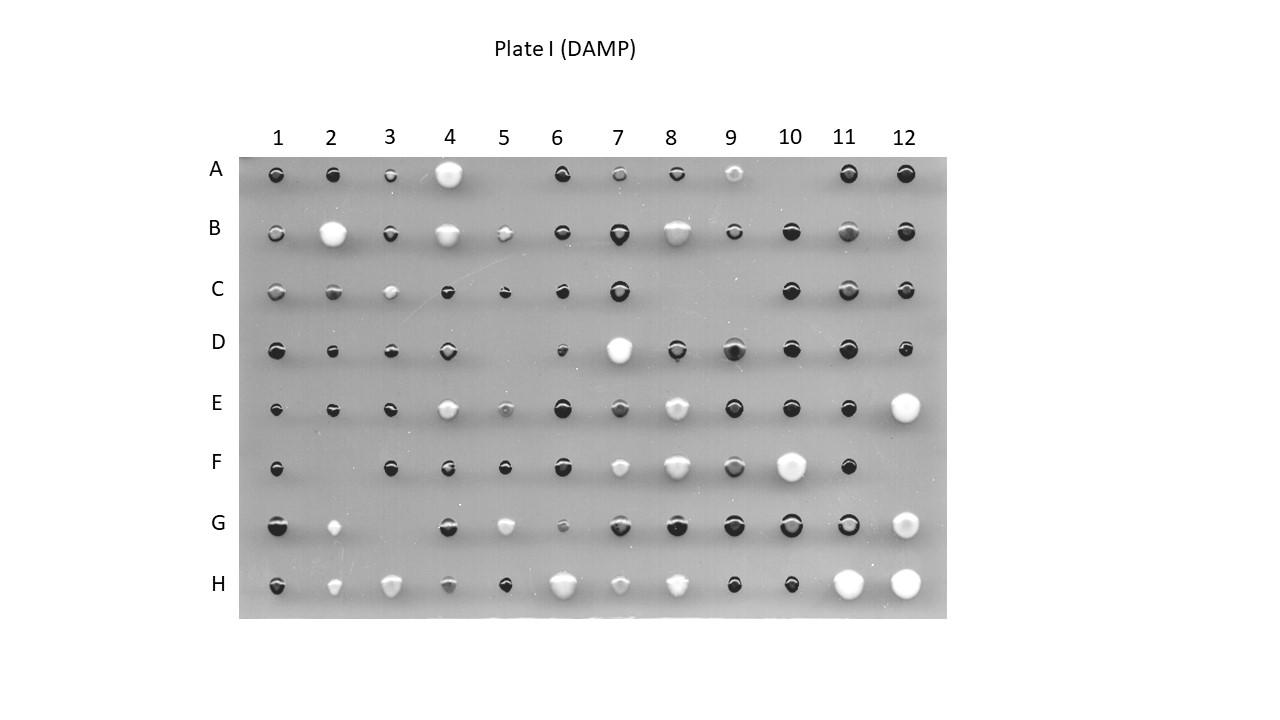

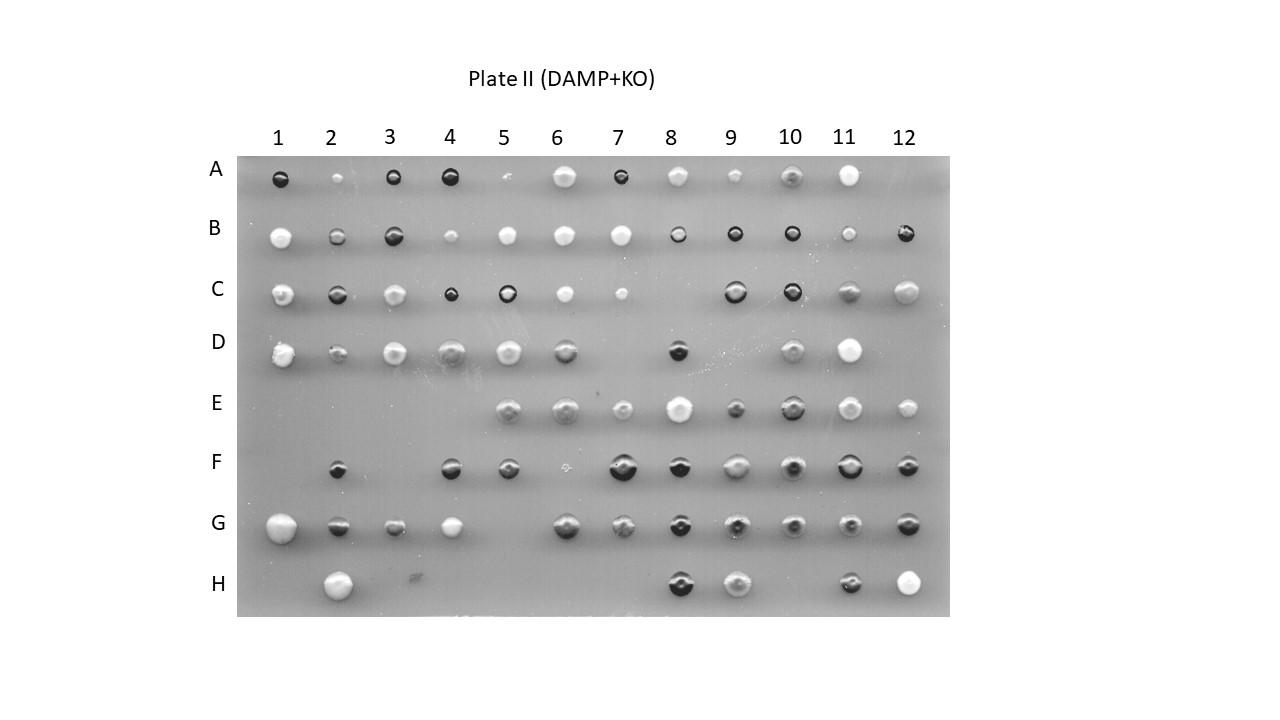

**Figure S1.** Plates of selected phloxine B positive mutants grown on YPD+phloxine B. Clones selected from the DAMP collection (Plate I) and from the DAMP and KO collection (Plate II). See Table S1 for plate maps.

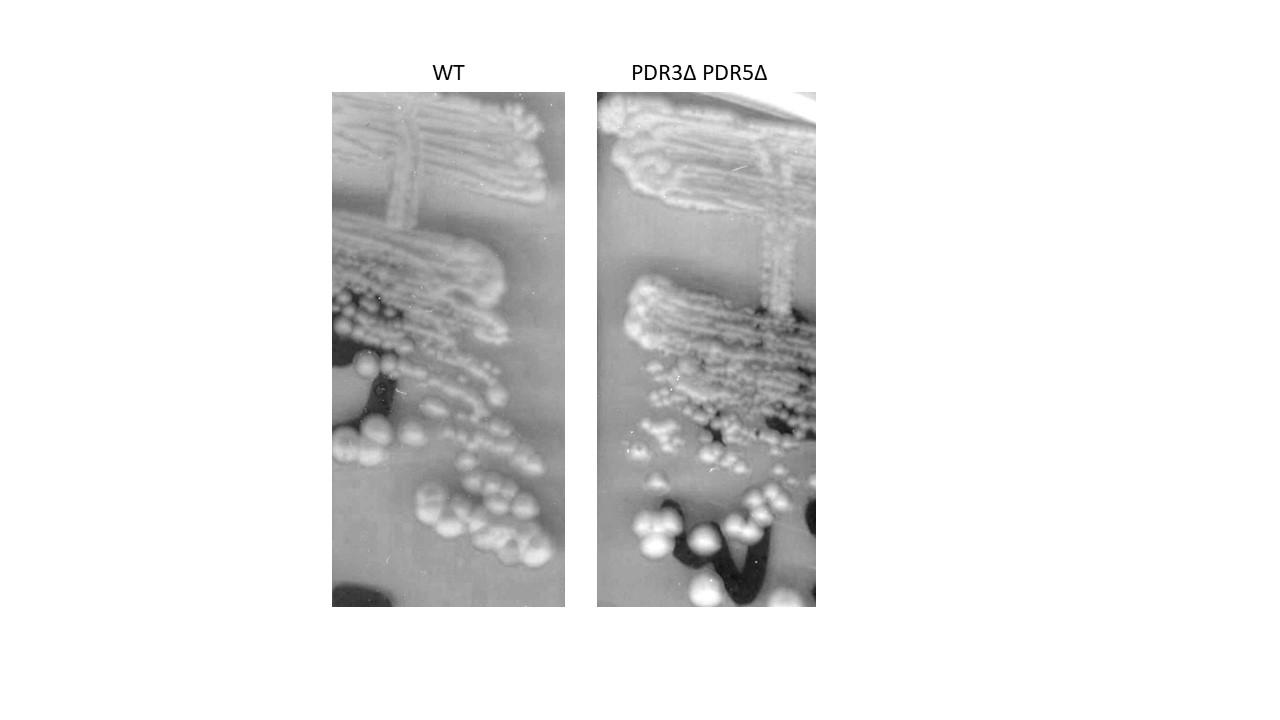

**Figure S2.** Deletion of drug efflux pump does not increase phloxine staining**.** The W303 strain (*leu2-3,112 trp1-1 can1-100 ura3-1 ade2-1 his3-11,15*) and its isogenic mutant strain with a deletion of the genes *PDR3* and *PDR5* were streaked onto a YPD plate containing phloxine B and grown for 48 hours.

**A**

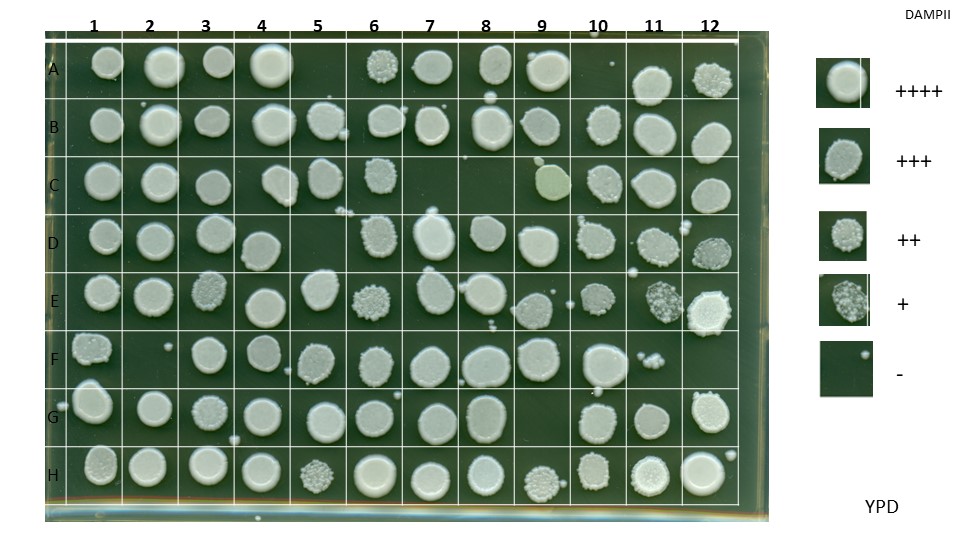

**B**

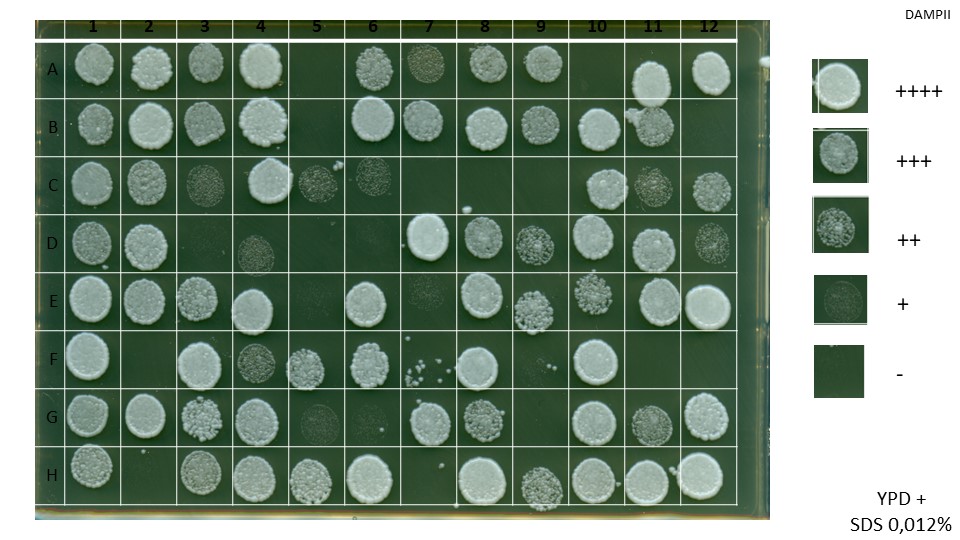

**C**

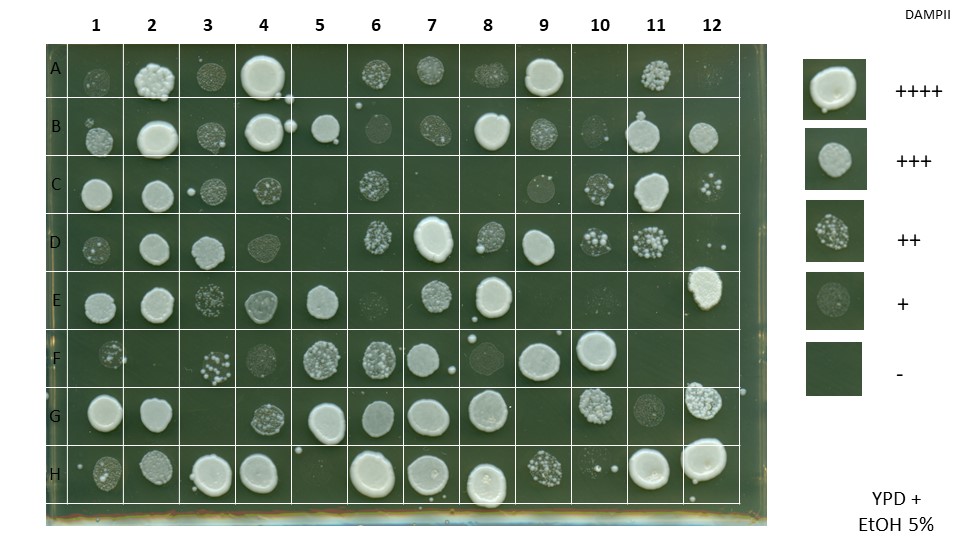

**D**

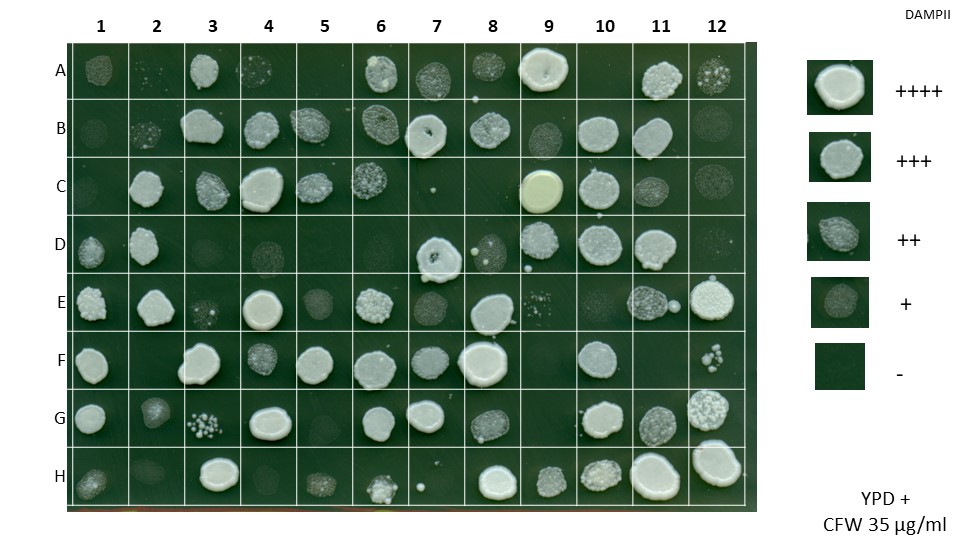

**E**

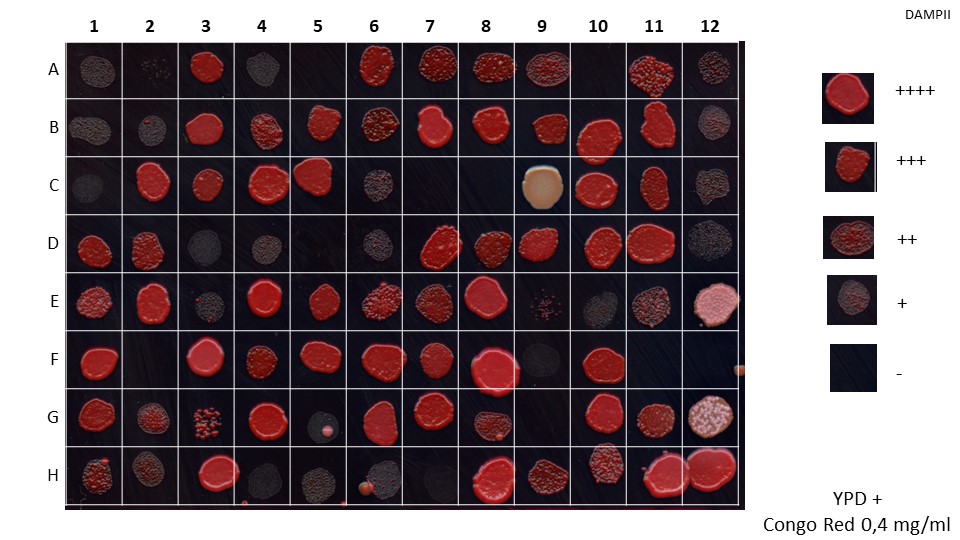

**F**

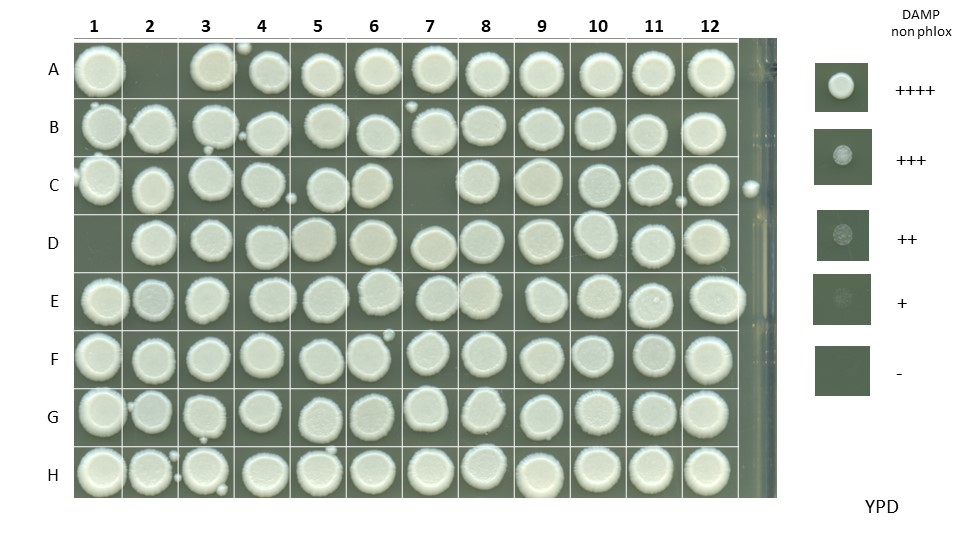

**G**
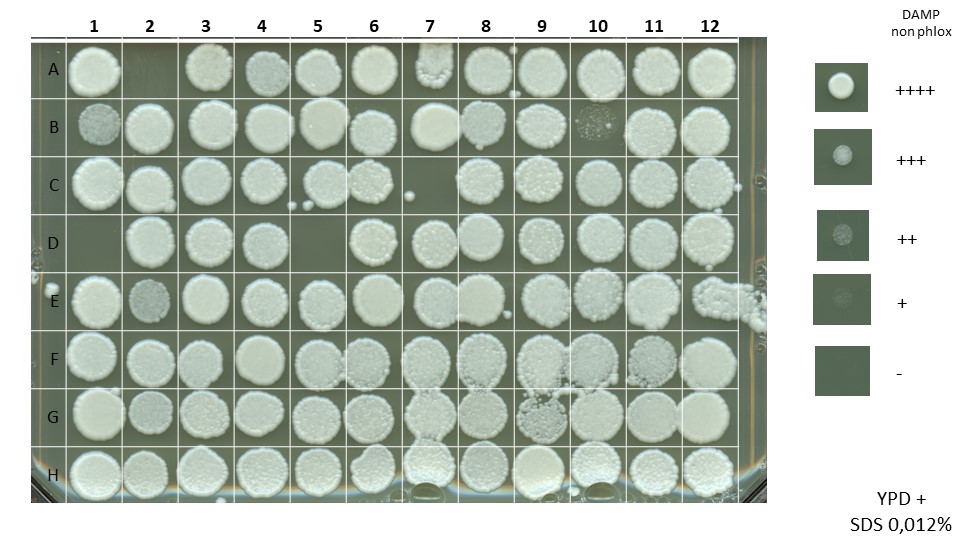

**H**
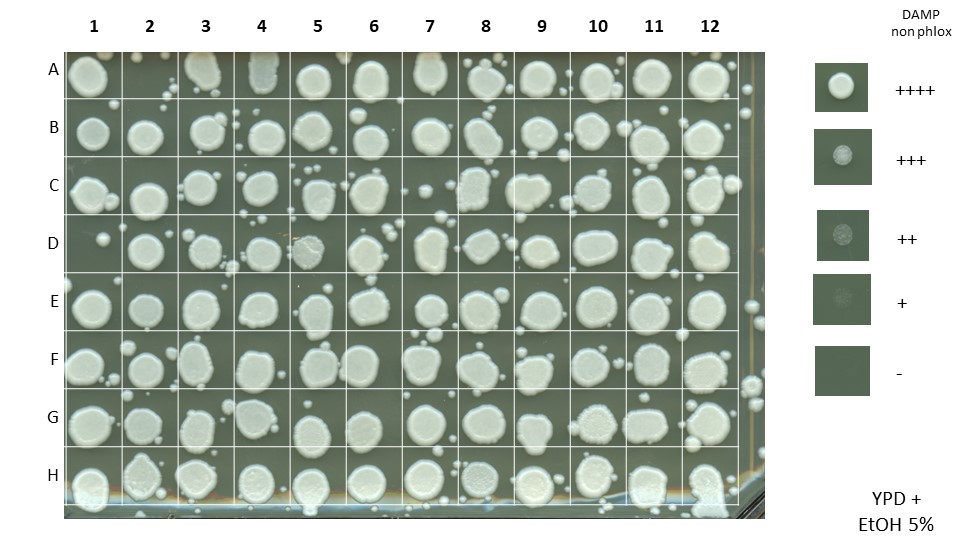

**I**
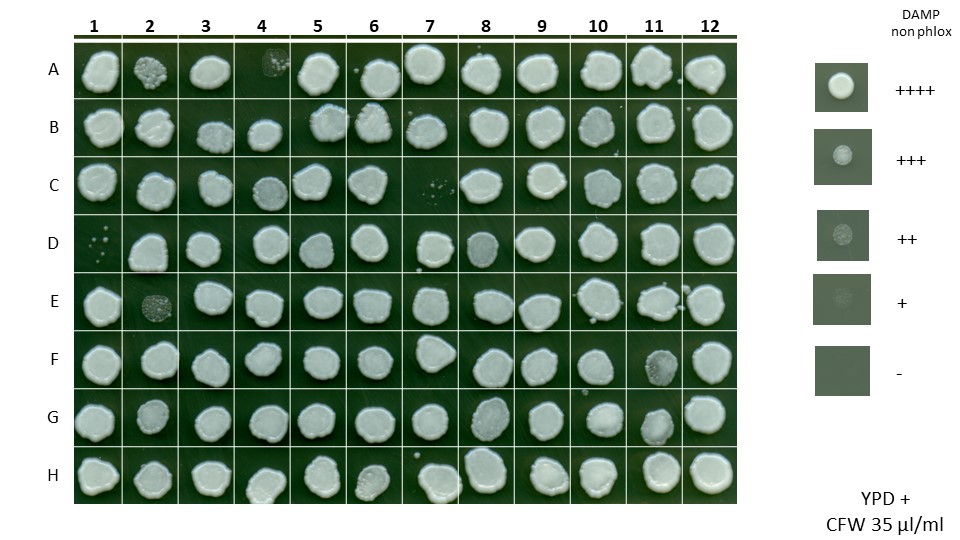

**J**
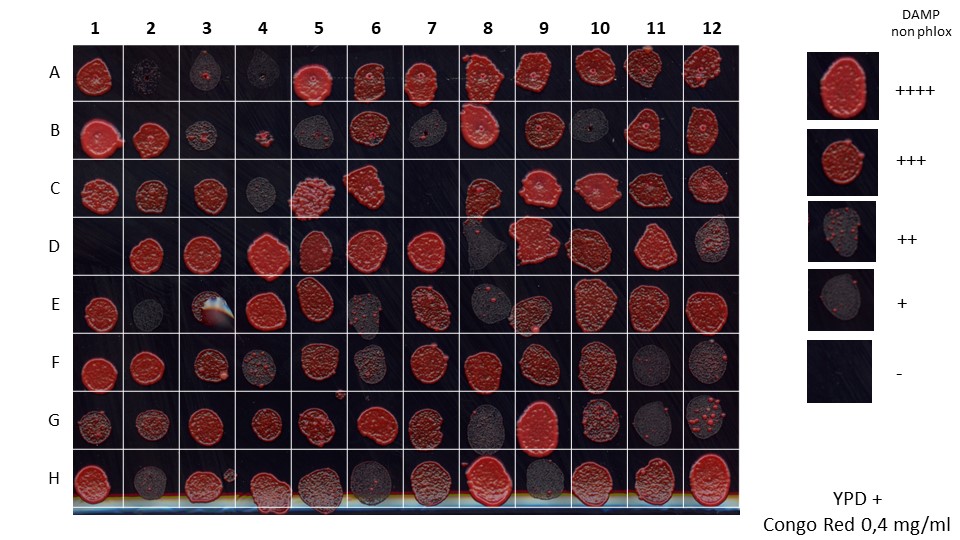

**Figure S3. Growth of select phloxine positive and phloxine negative mutants on media with various stressors.** (A) (B) (C) (D) (E) - test of growth of phloxine positive mutants; (F)(G)(H)(I)(J) - test of growth of phloxine negative mutants. H12 position is wild-type. (A)(F) YPD (B)(G) YPD+0,012% SDS (w.v) (C)(H) YPD+5% ethanol (v.v.) (D)(I) YPD+35mcg/ml CFW (E)(J) YPD+0,4 mg/ml Congo Red

**Figure S4. Various treatments reduce phloxine staining in different mutants.**

**A**

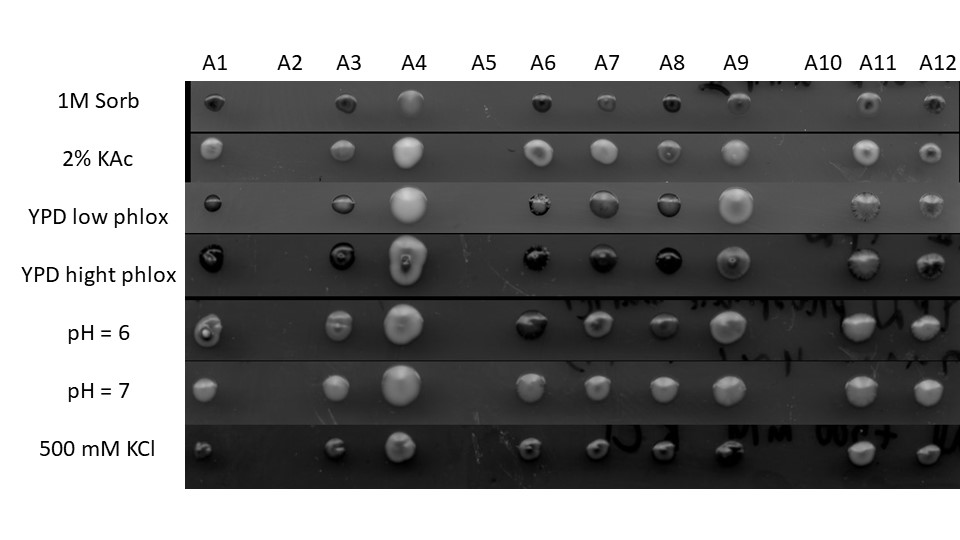

**B**

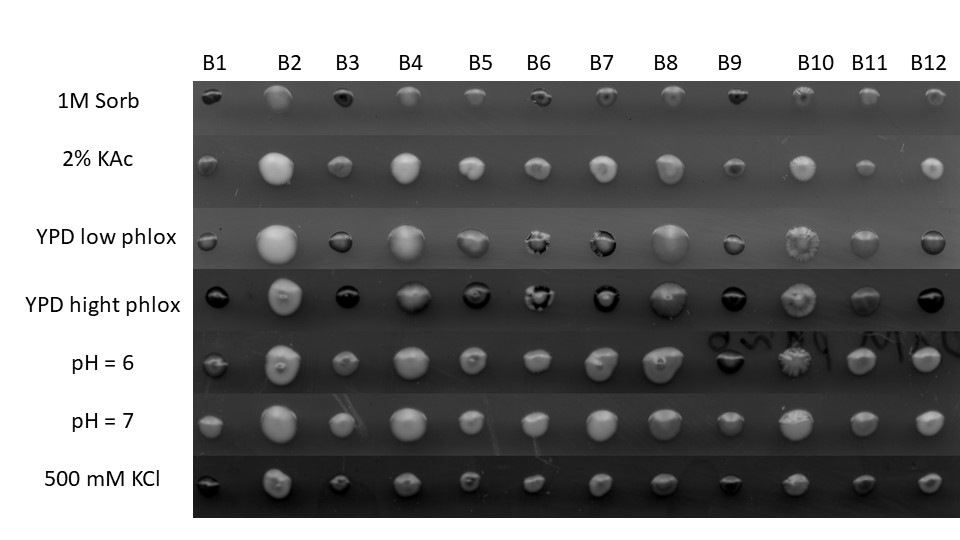

**C**

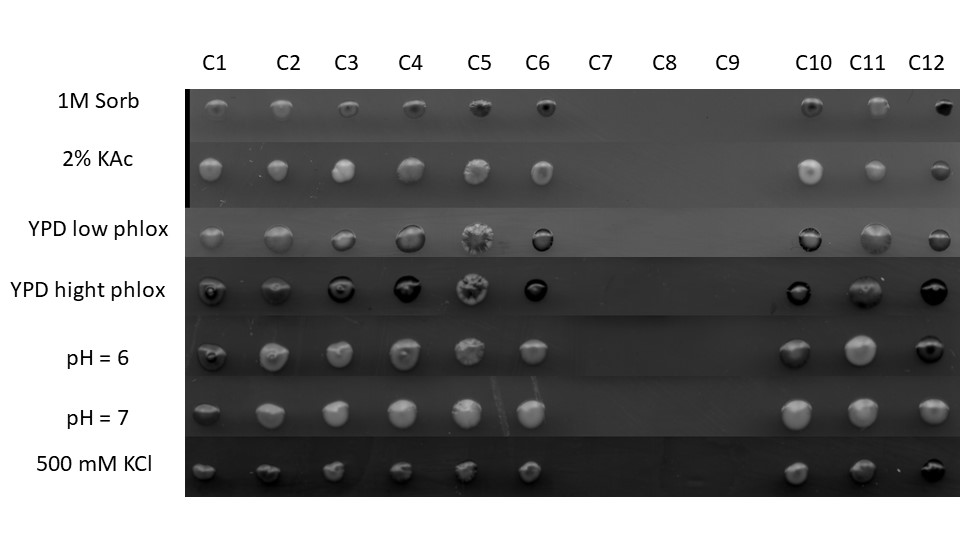

**D**
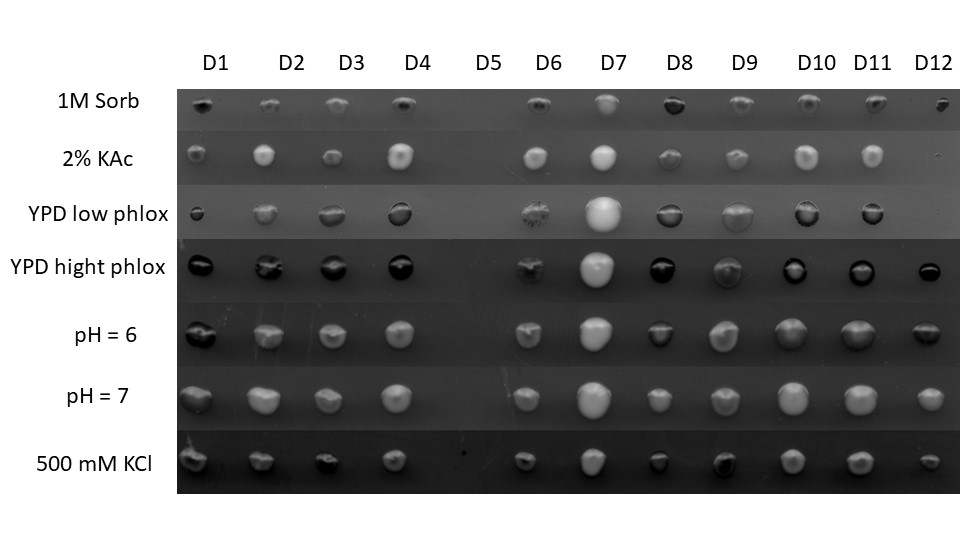

**E**
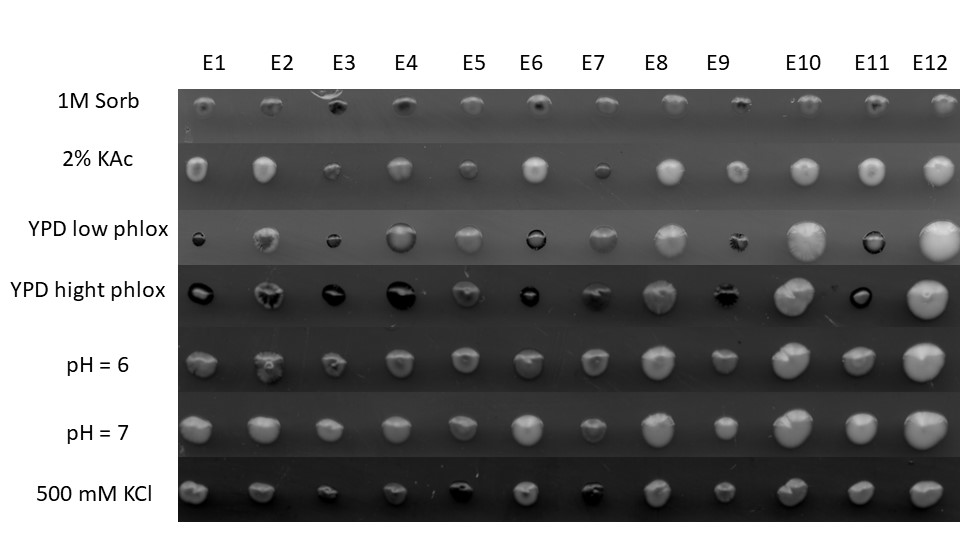

**F**

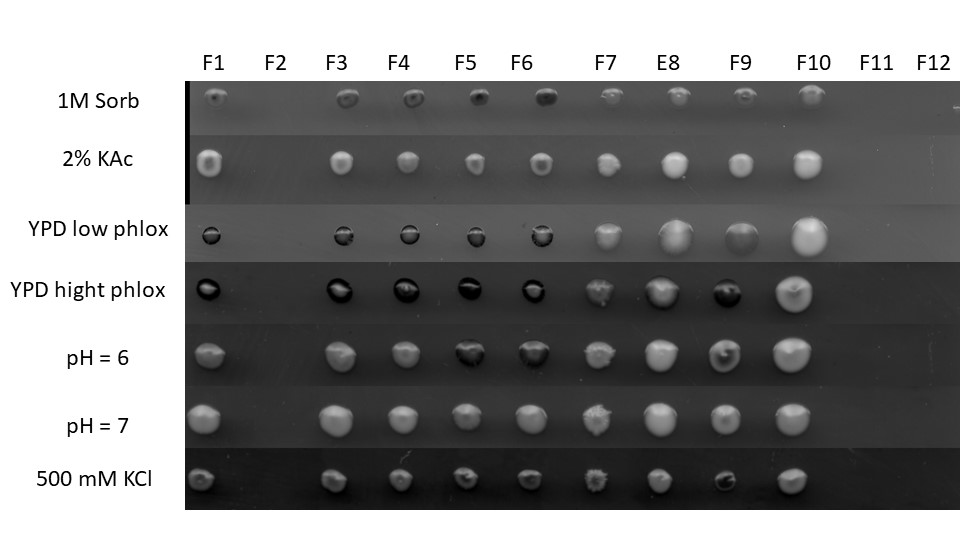

**G**

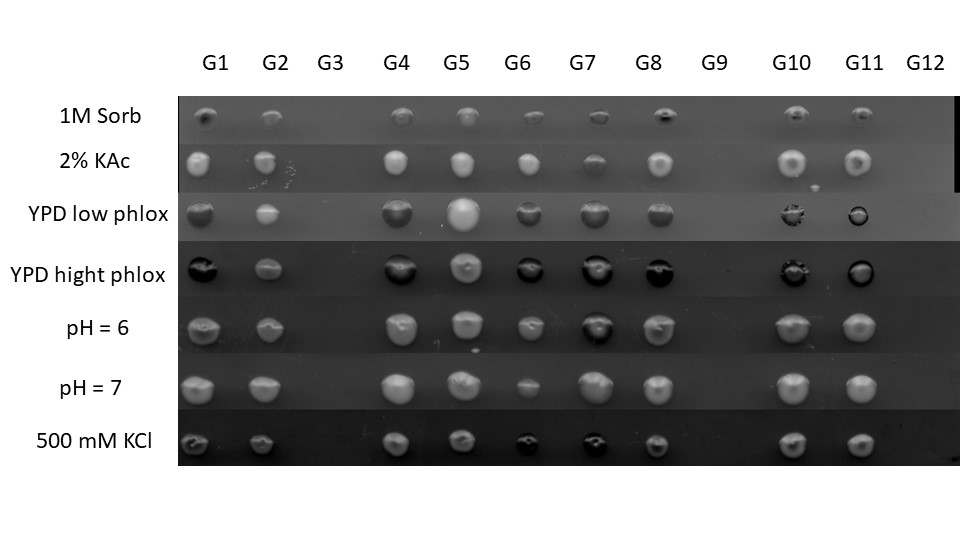

**H**

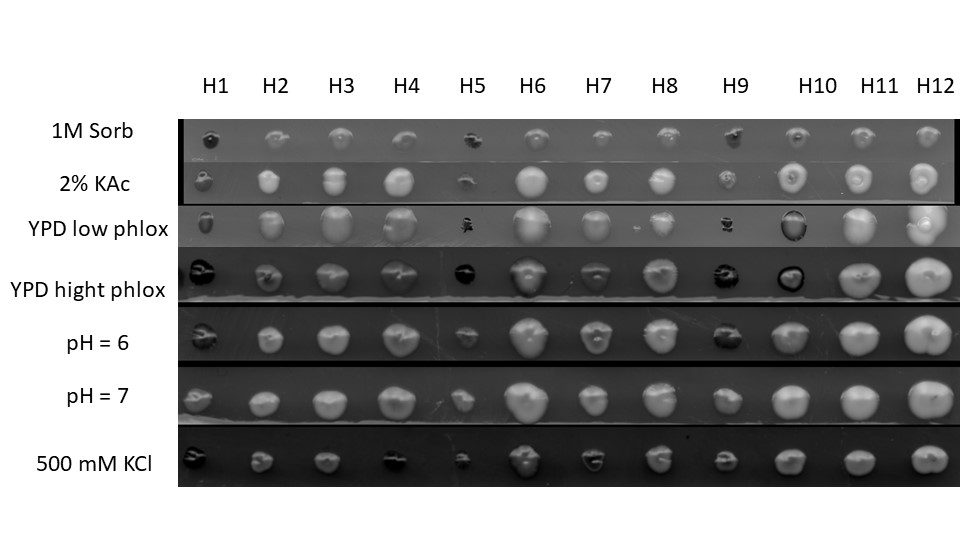

**Video 1**. Cells with downregulation of *CFT1* experiencing cell death during bud generation

**Video 2**. Cells with downregulation of *MSL5* experiencing death during cytokinesis

**Video 3**. Cells with downregulation of *MSL5* experiencing death during cytokinesis and demonstrating phloxine staining of a structure located at the bud neck.

**Video format note** – all the videos can be opened using VLC Media Player.
